# Loss of *Esr1* Does Not Affect Hearing and Balance

**DOI:** 10.1101/2024.03.03.583163

**Authors:** Shion S Simms, Marcus N Milani, Mi-Jung Kim, Ryan Husain, Laura Infante, Paul S Cooke, Shinichi Someya

## Abstract

Although estrogen affects the structure and function of the nervous system and brain and has a number of effects on cognition, its roles in the auditory and vestibular systems remain unclear. The actions of estrogen are mediated predominately through two classical nuclear estrogen receptors, estrogen receptor 1 (ESR1) and estrogen receptor 2 (ESR2). In the current study, we investigated the roles of ESR1 in normal auditory function and balance performance using 3-month-old wild-type (WT) and *Esr1* knockout (KO) mice on a CBA/CaJ background, a normal-hearing strain. As expected, body weight of *Esr1* KO females was lower than that of *Esr1* KO males. Body weight of *Esr1* KO females was higher than that of WT females, while there was no difference in body weight between WT and *Esr1* KO males. Similarly, head diameter was higher in *Esr1* KO vs. WT females. Contrary to our expectations, there were no differences in auditory brainstem response (ABR) thresholds, ABR waves I-V amplitudes and ABR waves I-V latencies at 8, 16, 32, and 48 kHz, distortion product otoacoustic emission (DPOAE) thresholds and amplitudes at 8, 16, and 32 kHz, and rotarod balance performance (latency to fall) between WT and *Esr1* KO mice. Furthermore, there were no sex differences in ABRs, DPOAEs, and rotarod balance performance in *Esr1* KO mice. Taken together, our findings show that *Esr1* deficiency does not affect auditory function or balance performance in normal hearing mice, and suggest that loss of *Esr1* is likely compensated by ESR2 or other estrogen receptors to maintain the structure and function of the auditory and vestibular systems under normal physiological conditions.

**Highlights:**

- Head diameter of female *Esr1* KO mice was higher than that of female WT mice.
- ABRs and DPOAEs were not different in WT and *Esr1* KO mice.
- There were no sex differences in ABRs and DPOAEs in *Esr1* KO mice.
- Rotarod balance performance was not different in WT and *Esr1* KO mice.
- There were no sex differences in rotarod balance performance in *Esr1* KO mice.
- Loss of *Esr1* does not affect auditory function or balance performance under normal physiological conditions.

## 1. Introduction

The gonadal steroid 17β-estradiol (E2) is a critical regulator of the male and female reproductive tract. However, actions of estrogen are not limited to the reproductive tract: estrogens also influence development and function of various non-reproductive tissues throughout the body, including the peripheral nervous systems (Behl et al., 2002; Charitidi et al., 2009; Williamson et al., 2019). Estrogen exerts the majority of its biological effects through interaction with estrogen receptor 1 and 2 (ESR1 and 2, respectively), both of which are found in the membrane and nucleus of target cells. In addition, the G protein-coupled receptor (GPER1) is exclusively localized in the cell membrane (Behl et al., 2002; Charitidi & Canlon, 2010; Charitidi et al., 2009) and is also expressed in many organs in the body. These estrogen receptors are expressed in brain regions and influence cognition and brain function. For example, estrogen is thought to influence memory through ESR1 and ESR2, which rapidly activate membrane estrogen signaling cascades involved in synaptic plasticity processes (Bean et al., 2014). In mice, the absence of the *Esr1* gene impairs spatial discrimination (Fugger et al., 1998), while lentiviral delivery of *Esr1* into the hippocampus of *Esr1* KO mice restores spatial learning (Foster et al., 2008). In humans, ESR1 polymorphisms are associated with an increased risk of cognitive dysfunction (Olsen et al., 2006; Yaffe et al., 2002).

A growing body of literature suggests that estrogen also influences auditory function. A cross-sectional study in postmenopausal women found a significant association between hearing sensitivity and serum estradiol levels (Kim et al., 2002). In pre-menopausal women, auditory sensitivity fluctuates during the menstrual cycle with the lowest hearing thresholds, or heightened hearing sensitivity, occurring during the late-follicular phase of the menstrual cycle, which corresponds to the highest levels of serum estrogen (Souza et al., 2017). Interestingly, estrogen replacement therapy lowers hearing thresholds, shortens auditory brainstem response (ABR) latencies, and increases ABR amplitudes in postmenopausal women (Hederstierna et al., 2007; Khaliq et al., 2005; Khaliq et al., 2003). In line with these reports, Turner syndrome, a condition that results when one of the X chromosomes is completely or partially missing in women, is commonly associated with ovarian insufficiency, estrogen deficiency, and early onset sensorineural hearing loss (Hederstierna et al., 2009). In rats, ovariectomized females display longer ABR latencies compared to controls, while replacement therapy with E2, the most potent endogenous form of estrogen, shortens ABR latencies (Coleman et al., 1994). In mice, E2-treated animals display lower ABR thresholds and higher ABR wave amplitudes compared to other hormone treatment subject groups (Williamson et al., 2019).

Both ESR1 and ESR2 are expressed in the inner ears of humans and mice (Motohashi et al., 2010; Stenberg et al., 1999). In the cochlea of rodents, ESR1 and ESR2 are expressed in the inner hair cells, outer hair cells, and spiral ganglion neurons (Stenberg et al., 1999). ESR1 is also present in the stria vascularis, spiral ligament, pillar, and phalangeal cells (Charitidi et al., 2009). However, a previous study has shown that the mean ABR threshold from young *Esr1*^-/-^ mice (*Esr1* KO mice) in the 129/J-C57BL/6J background does not differ from WT mice (Meltser et al., 2008), indicating that loss of *Esr1* has no direct effect on hearing function in 129/J-C57BL/6J mice.

In the current study, we investigated the roles of ESR1 in normal auditory function and balance performance using 3-month-old *Esr1*^+/+^ (WT) and *Esr1*^-/-^ (*Esr1* KO) mice on a CBA/CaJ background, a normal-hearing strain. We used CBA/CaJ mice because even though most studies looking at estrogen effects have utilized *Esr1* KO mice on a C57BL/6J background, this strain is homozygous for the age-related hearing loss (*ahl*)-susceptibility allele (*Cdh23*^753A^) and displays early onset age-related hearing loss (ARHL) by 9-12 months of age (Noben-Trauth et al., 2003; Zheng et al., 1999). Thus, examining how *Esr1* deficiency affects hearing sensitivity and balance performance is most effectively done with the CBA/CaJ mouse strain, which is a normal-hearing mouse and model for late-onset ARHL (Han and Someya, 2013; Johnson et al., 2006; Noben-Trauth et al., 2003).

## 2. Methods

### 2.1. Animals

Heterozygous *Esr1^+/-^* mice were a gift from Dr. Paul Cooke (University of Florida). Generation of *Esr1^-/-^* (*Esr1* KO) mice has been previously described (Dupont et al., 2000). CBA/CaJ mice were purchased from the Jackson Laboratory (https://www.jax.org/strain/000654). Both male and female mice were used in the current study. All animal experiments were conducted under protocols approved by the University of Florida Institutional Animal Care and Use Committee.

### 2.2. Backcrossing and genotyping

The *Esr1* mutant mice were originally generated on a mixed 129/Sv-C57BL/6J background (Dupont et al., 2000). Because the C57BL/6J mouse strain is homozygous for the age-related hearing loss (*ahl*)-susceptibility allele (*Cdh23*^753A^) (Noben-Trauth et al., 2003; Zheng et al., 1999), we removed the *ahl* allele by backcrossing *Esr1*^+/−^ mice onto the CBA/CaJ mouse strain, a normal-hearing strain homozygous for the AHL-resistance allele (*Cdh23*^753G^) for five generations (Park et al., 2019). To confirm that WT and *Esr1* KO mice have the same wild-type *Cdh23* genotype (*Cdh23*^753G/753G^), tail DNA from these mice was amplified by PCR, and then we sequenced the region of DNA containing the 753rd nucleotide in the *Cdh23* gene. Primer sequences and cycling conditions for PCR were as follows:

*Cdh23* forward 5’-GATCAAGACAAGACCAGACCTCTGTC-3’;
*Cdh23* reverse 5’-GAGCTACCAGGAACAGCTTGGGCCTG-3’;

PCR cycling conditions consisted of an initial denaturation at 95 °C for 2 min, 35 cycles of 95 °C for 30 sec, 60 °C for 1 min, 72 °C for 1 min, and 72 °C for 5 min. The expected size of the PCR product was 360 bps. All the mice used in this study had the same wild-type *Cdh23* genotype (*Cdh23*^753G/753G^).

CBA/CaJ-*Esr1*^+/-^ males were then mated with CBA/CaJ-*Esr1*^+/-^ females and their offspring were genotyped via PCR using DNA isolated from the tails of the weaned mice at 3 weeks of age. Primer sequences and cycling conditions for PCR were as follows:

*Esr1* forward 5’ - TCG CTT TCC TGA AGA CCT TCC ATA T - 3’;
*Esr*1 reverse 5’ - CCA TTG TCT CTT TCT GAC ACA TGC - 3’;
*Esr*1 KO reverse 5’ - GCA AAT AGC GGG AGA TCT AAT TCT AGA TAA C - 3’;

PCR cycling conditions consisted of an initial denaturation at 94 °C for 5 min, followed by denaturation at 94 °C for 30 sec, annealing at 55 °C for 30 sec, and extension at 72 °C for 90 sec. 34 cycles of 94 °C for 30 sec were repeated, followed by a final soak at 72 °C for 5 min. PCR products were then loaded onto a 2% agarose gel and electrophoresed for 25 min at 100 V. The expected band sizes for the WT and KO alleles were 280 and 190 base pairs, respectively (Fig. 1).

**Fig. 1.**
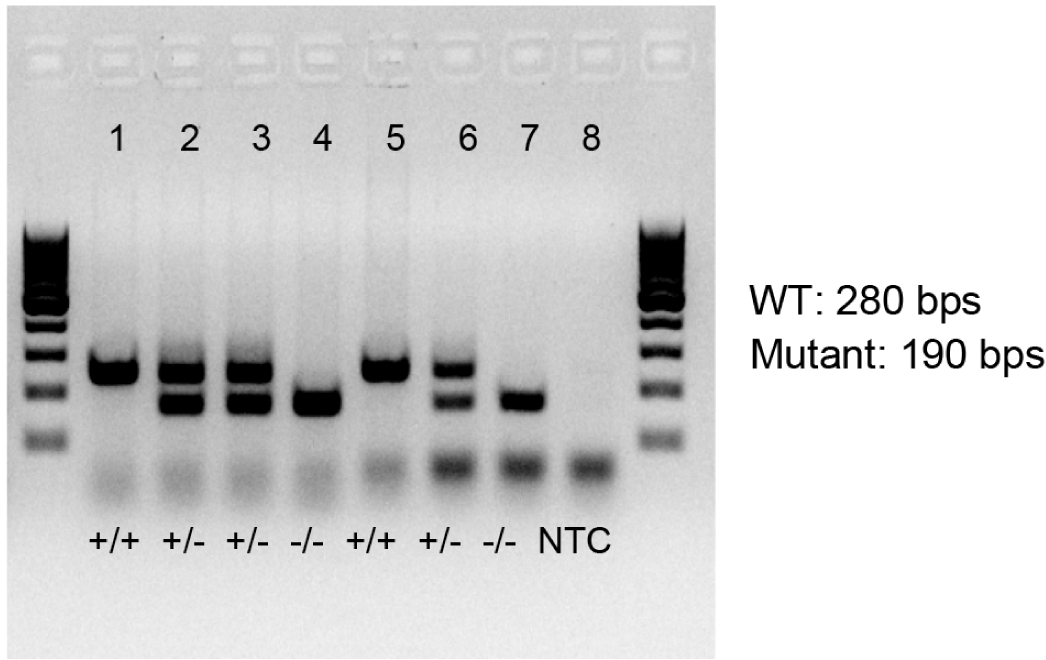
Genotyping. *Esr1* genotyping: PCR products were separated on a 2% agarose gel. The expected band sizes for the wild-type and mutant alleles were 280 and 190 bps, respectively. +/+, *Esr1*^+/+^; +/-, *Esr1^+/-^*; -/-, *Esr1*^-/-^; WT, wild-type; NTC, no template control.

### 2.3. Bodyweight and head diameter

Body weights of WT and *Esr1* KO mice (10 WT males, 9 *Esr1* KO males, 10 WT females, and 8 *Esr1* KO females) were measured prior to ABR testing at 3 months of age. Head diameter of WT and *Esr1* KO mice (7 WT males, 4 *Esr1* KO males, 8 WT females, and 5 *Esr1* KO females) was measured following DPOAE testing at 3 months of age.

### 2.4. Auditory brainstem response

ABR hearing tests were conducted using an ABR and DPOAE recording system with RZ6 hardware and BioSigRZ software (Tucker-Davis Technologies), as previously described (Kim et al., 2023; Kim et al., 2020; Kim et al., 2019). Mice were anesthetized via intraperitoneal injection (ip) of ketamine (100 mg/kg) and xylazine (10 mg/kg). Subdermal needle electrodes were positioned at the vertex (active), ipsilateral ear (reference), and contralateral ear (ground). Sound output from the speaker was directed through a PVC tube into the right ear canal of the animals (closed field). Tone calibration was performed according to the guidelines provided in the TDT ABR User Guide (https://www.tdt.com/files/manuals/ABRGuide.pdf).

ABR thresholds were assessed using tone burst stimuli at 8, 16, 32, and 48 kHz. At each frequency, sound intensity was incrementally reduced in 10 dB sound pressure level (SPL) steps, ranging from 90 to 10 dB SPL. A threshold was identified as the lowest stimulus level at which response peaks for wave I were visibly present upon visual inspection. ABR amplitudes and latencies for ABR waves I, II, III, IV, and V were measured using an 80 dB SPL tone burst stimulus at 8, 16, 32, and 48 kHz. A wave amplitude was determined by measuring the voltage difference between the peak and trough of each ABR wave (Kim et al., 2023). Wave latency was calculated by measuring the time elapsed from the stimulus onset to the peak of each ABR wave (Kim et al., 2023). We used 10 WT males, 9 *Esr1* KO males, 10 WT females, and 8 *Esr1* KO females at 3 months of age for ABR threshold, wave amplitude, and wave latency measurements.

### 2.5. Distortion Product Otoacoustic Emissions (DPOAE)

Three days following the ABR tests, DPOAE tests were conducted using the ABR and DPOAE recording system with RZ6 hardware and BioSigRZ software (Tucker-Davis Technologies) according to TDT’s DPOAE User Guide (https://www.tdt.com/files/manuals/DPOAEGuide.pdf) and as previously described (Manohar et al., 2022). Mice were anesthetized through ip injection of ketamine and xylazine, as described above. The tip connected to the ER10B+ (Etymotic Research) and two MF1 speakers (Tucker-Davis Technologies) was positioned in the right ear canal. DPOAE amplitudes at 2F1-F2 were collected in response to two-tone sound stimuli. The tone frequencies F1 and F2 had a F2/F1 ratio of 1.2 and were geometrically centered about 8, 16, and 32 kHz. At each center frequency, the tone levels L1 and L2 remained equal and were reduced in 10LdB SPL steps from 80 to 20LdB SPL. A DPOAE threshold was defined as the lowest stimulus level at which the 2F1-F2 distortion product was more than 6 dB SPL above the noise floor (Powers et al., 2006). A noise floor was defined as the average of the distortion products at ten neighboring frequencies (five above and five below the 2F1-F2) (Powers et al., 2006). We used 10 WT males, 9 *Esr1* KO males, 10 WT females, and 8 *Esr1* KO females at 3 months of age for DPOAE amplitude and threshold measurements.

### 2.6. Rotarod balance performance

Prior to ABR testing, mice were trained to use a rotarod apparatus (Rota Rod Rotamex 5; Columbus Instruments) with an acceleration of 2 rpm every 17 sec starting from 4 rpm to 40 rpm during the first five trials (trials 1-5). The rotarod balance performance (average latency to fall) was recorded during the next three trials (trials 6-8) as previously described (Kim et al., 2023). During training trials 1-5, mice were tested for ∼90 sec, with a 1 min rest between each trial. In the subsequent tested trials 6-8, mice were tested for ∼5 min, with a 3 min rest between each trial. We used 10 WT males, 9 *Esr1* KO males, 10 WT females, and 8 *Esr1* KO females at 3 months of age.

### 2.7. Statistical analysis

Two-way ANOVA with Bonferroni post hoc testing (GraphPad Prism 10) was used to analyze the ABR thresholds and DPOAE thresholds. One-way ANOVA with Tukey’s multiple comparisons tests post-hoc testing (GraphPad Prism 10) was used to analyze ABR wave I-V amplitudes and wave I-V latencies, DPOAE amplitudes, bodyweight, head diameters, and latencies to fall off the rotarod.

## 3. Results

### 3.1. Effects of Esr1 deficiency on body weight and head diameter

The full disruption of the gene for ESR1 was confirmed by the absence in the *Esr1* KO mice, and no *Esr1* mRNA was detected by RT-PCR as previously described (Dupont et al., 2000). Because the *Esr1* mutant mice were originally generated in the mixed 129/Sv-C57BL/6J background (Dupont et al., 2000) but C57BL/6J mice have an age-related hearing loss (*ahl*)-susceptibility allele (*Cdh23*^753A^) (Noben-Trauth et al., 2003; Zheng et al., 1999), for our experiments we used mice that were backcrossed *Esr1*^+/−^ mice onto the CBA/CaJ mouse strain, a normal-hearing strain homozygous for the AHL-resistance allele (*Cdh23*^753G^) for five generations (Park et al., 2019). Homozygous CBA/CaJ mutant mice lacking *Esr1* appeared to develop normally.

It has been suggested that the amplitude and latency of ABR waves can be affected by many factors such as body size and head size (Zhou et al., 2006), and that small head size and or a shorter cochlear length in females may contribute to greater synchronous activity at the afferent auditory pathway, leading to greater ABR wave amplitudes and shorter wave latencies (McFadden, 1998; Shuster et al., 2019). To examine how *Esr1* deficiency affects body and head size in *Esr1* KO mice, we measured body weight and head diameter of WT and *Esr1* KO females and males at 3 months of age. As expected, body weight of WT females was lower than that of WT males (Fig. 2A). Similarly, body weight of *Esr1* KO females was lower than that of *Esr1* KO males. Body weight of *Esr1* KO females was higher than that of WT females, while body weights in WT and *Esr1* KO males did not differ. Consistent with the body weights, head diameter of WT females was lower than that of WT males (Fig. 2B), while head diameter of *Esr1* KO females was higher than that of WT females. There was no difference in head diameter between *Esr1* KO males and *Esr1* KO females.

**Fig. 2.**
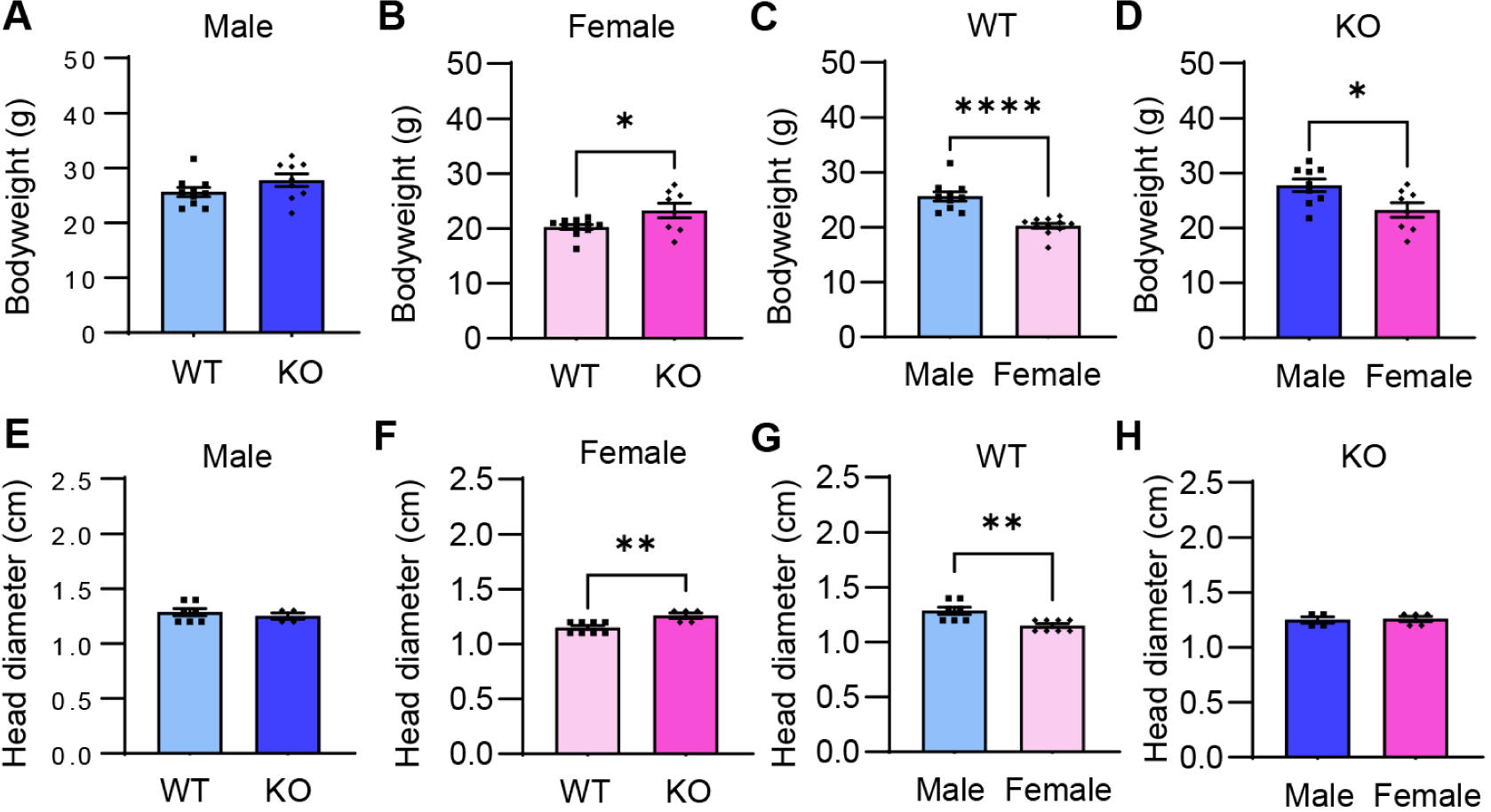
Bodyweight and head diameter. (A) Body weight was measured in WT and *Esr1* KO mice at 3 months of age; male WT (n=10), male KO (n=9), female WT (n=10), female KO (n=8). (B) Head diameter was measured in WT and *Esr1* KO mice at 3 months of age; male WT (n=7), male KO (n=4), female WT (n=8), female KO (n=5). Data are shown as means ± SEM. One-way ANOVA with Tukey’s multiple comparisons tests was performed. *, *p*<0.05; **, *p*<0.01; ****, *p*<0.0001.

### 3.2. Effects of Esr1 deficiency on auditory function

To examine how *Esr1* deficiency affects hearing sensitivity, we measured ABR thresholds, waves I-V amplitudes, and waves I-V latencies at 8, 16, 32, and 48 kHz, and DPOAE thresholds and amplitudes at 8, 16, and 32 kHz in WT and *Esr1* KO mice at 3 months of age. Contrary to our expectations, there were no differences in ABR thresholds (Fig. 3A-B), ABR waves I-V amplitudes (Fig. 4A-H), and ABR waves I-V latencies (Fig. 5A-H) at all the frequencies measured between WT and *Esr1* KO males or WT and *Esr1* KO females. There were no sex differences in ABR thresholds (Fig. 3C-D), waves I-V amplitudes (Fig. 4I-P), and waves I-V latencies (Fig. 5I-P) in WT or *Esr1* KO mice. In agreement with the ABR test results, there were no differences in DPOAE amplitudes (Fig. 6A-F) and thresholds (Fig. 7A-B) at all the frequencies measured between WT and *Esr1* KO males or WT and *Esr1* KO females, and there were no sex differences in DPOAE amplitudes (Fig. 6G-L) and thresholds (Fig. 7C-D) in WT or *Esr1* KO mice.

**Fig. 3.**
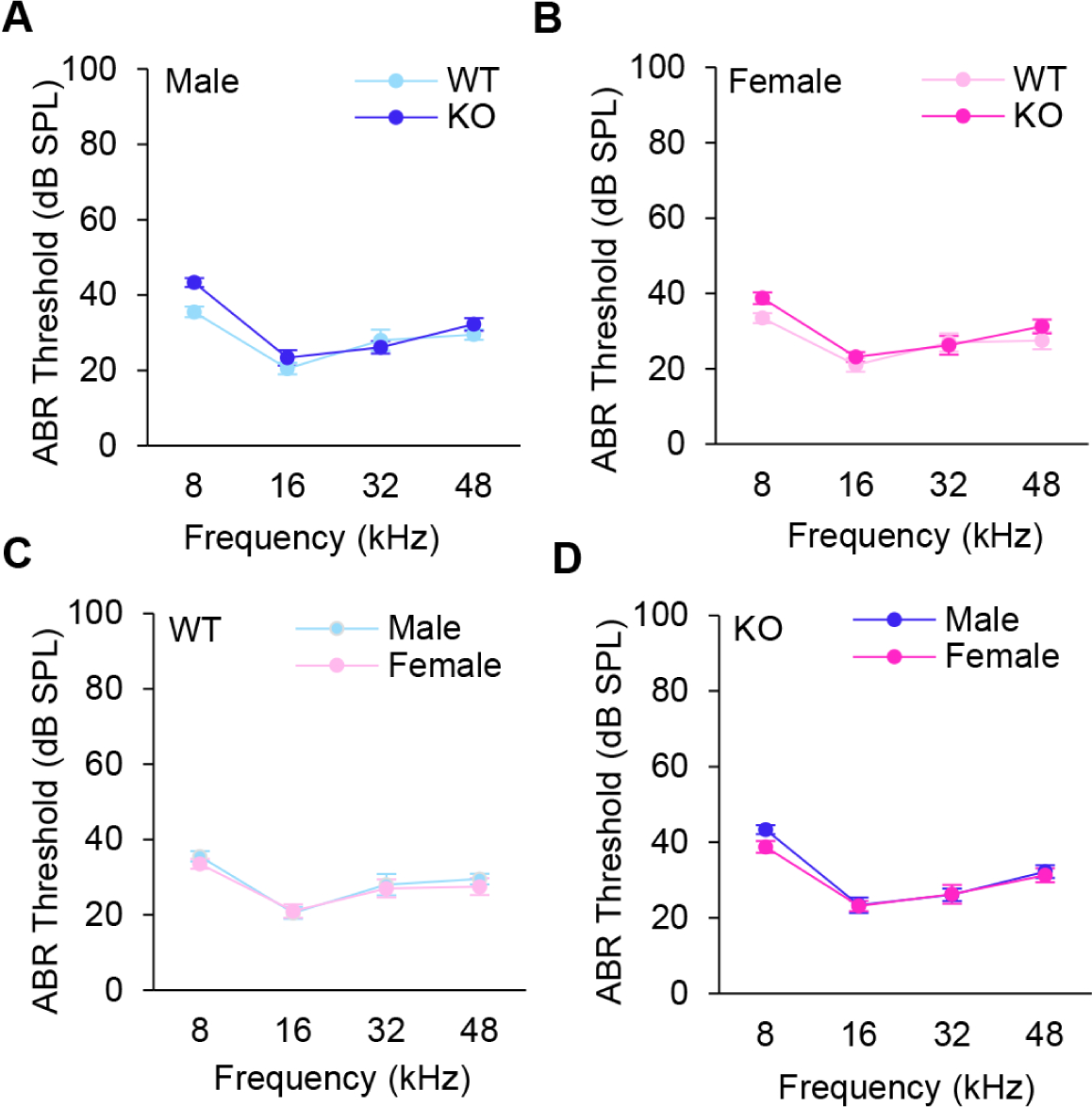
ABR threshold. (A-D) ABR thresholds were measured with tone burst stimuli at 8, 16, 32, and 48 kHz in WT and *Esr1* KO mice at 3 months of age; male WT (n=10), male KO (n=9), female WT (n=10), female KO (n=8). Data are shown as means ± SEM; Two-way ANOVA with Bonferroni’s multiple comparisons tests were performed. *p*<0.05.

**Fig. 4.**
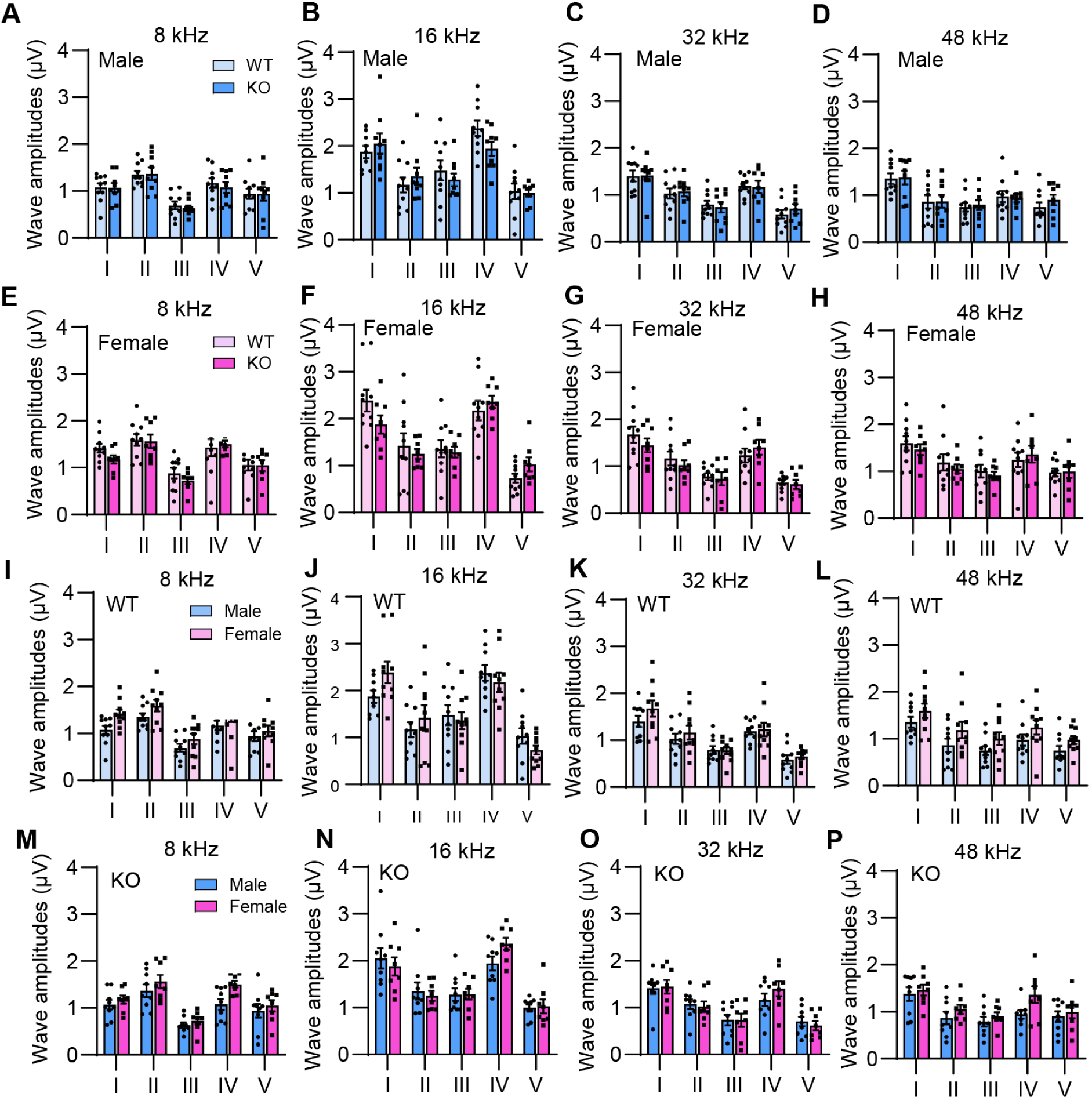
ABR Waves I-V Amplitude. (A-P) ABR Waves I-V amplitudes were measured at 8, 16, 32, and 48 kHz at 80 dB SPL in WT and *Esr1* KO mice at 3 months of age; male WT (n=10), male KO (n=9), female WT (n=10), female KO (n=8). Data are shown as means ± SEM; One-way ANOVA with Tukey’s multiple comparisons tests was performed. *p*<0.05.

**Fig. 5.**
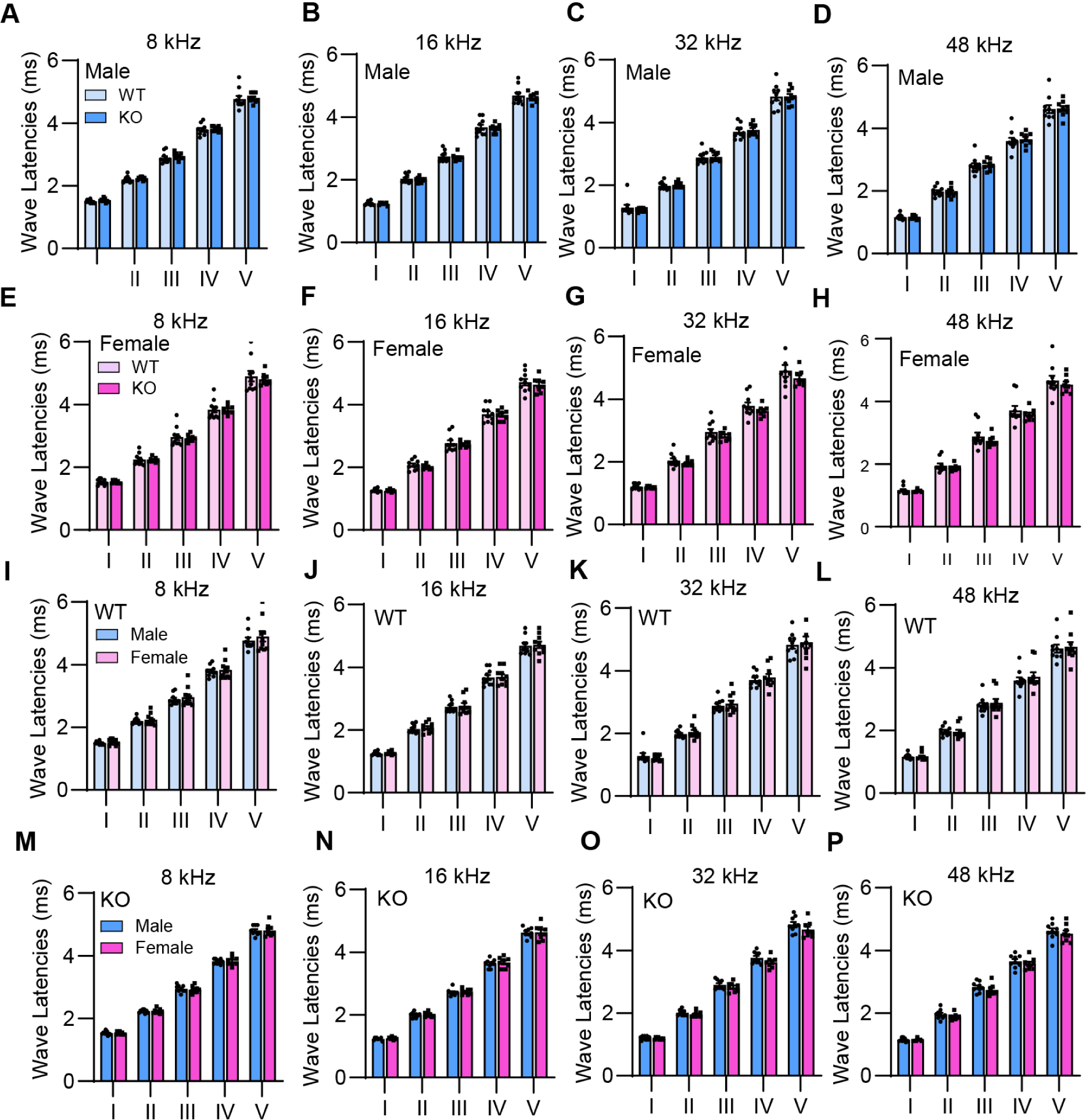
ABR Waves I-V Latency. (A-P) ABR Wave I-V latencies were measured at 8, 16, 32, and 48 kHz at 80 dB SPL in WT and *Esr1* KO mice at 3 months of age; male WT (n=10), male KO (n=9), female WT (n=10), female KO (n=8). Data are shown as means ± SEM; One-way ANOVA with Tukey’s multiple comparisons tests was performed. *p*<0.05.

**Fig. 6.**
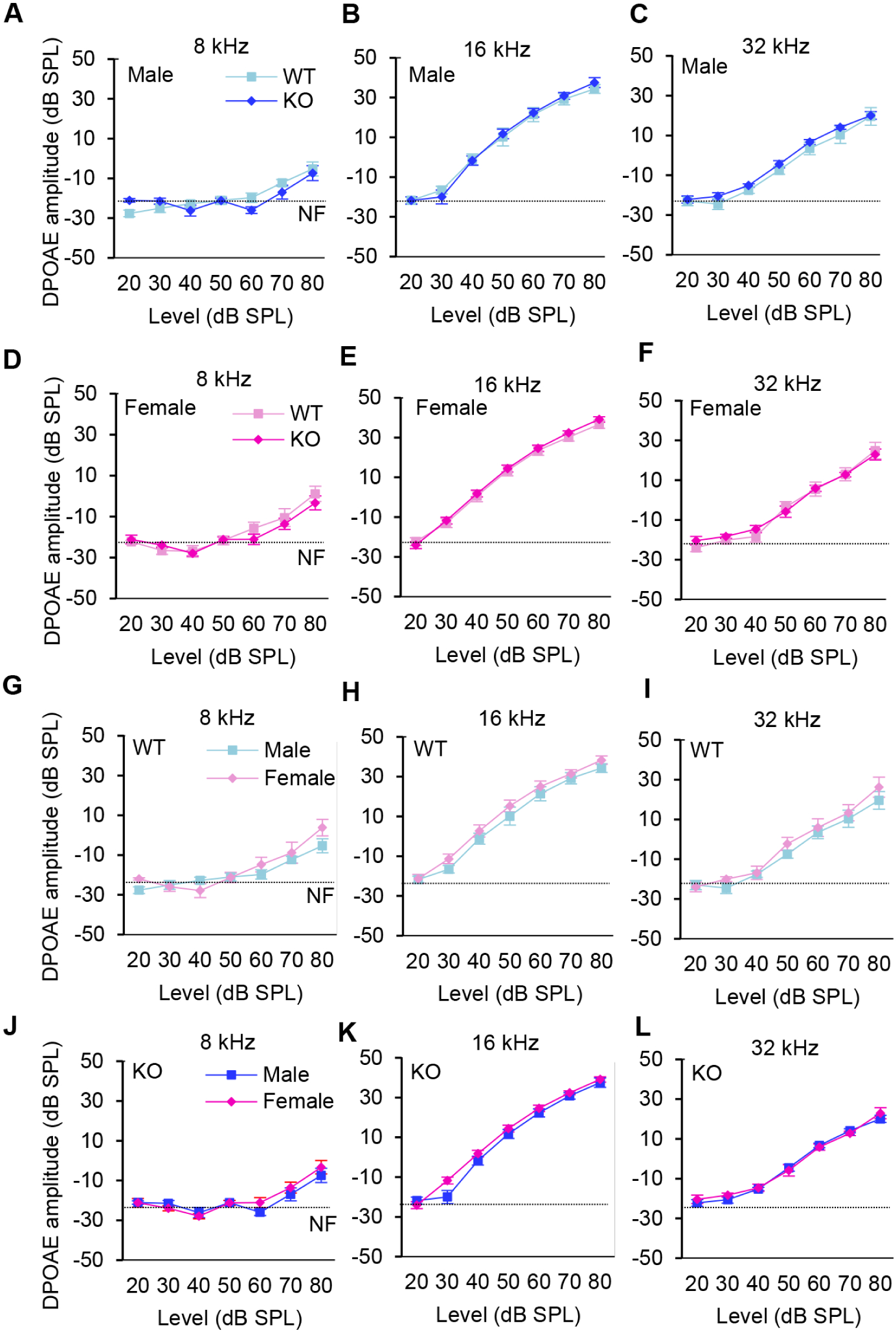
DPOAE Amplitude. (A-L) DPOAE amplitudes were measured at 8, 16, and 32 kHz in WT and *Esr1* KO mice at 3 months of age; male WT (n=10), male KO (n=9), female WT (n=10), female KO (n=8). Data are shown as means ± SEM. One-way ANOVA with Tukey’s multiple comparisons tests was performed. *p*<0.05. NF, noise floor.

**Fig. 7.**
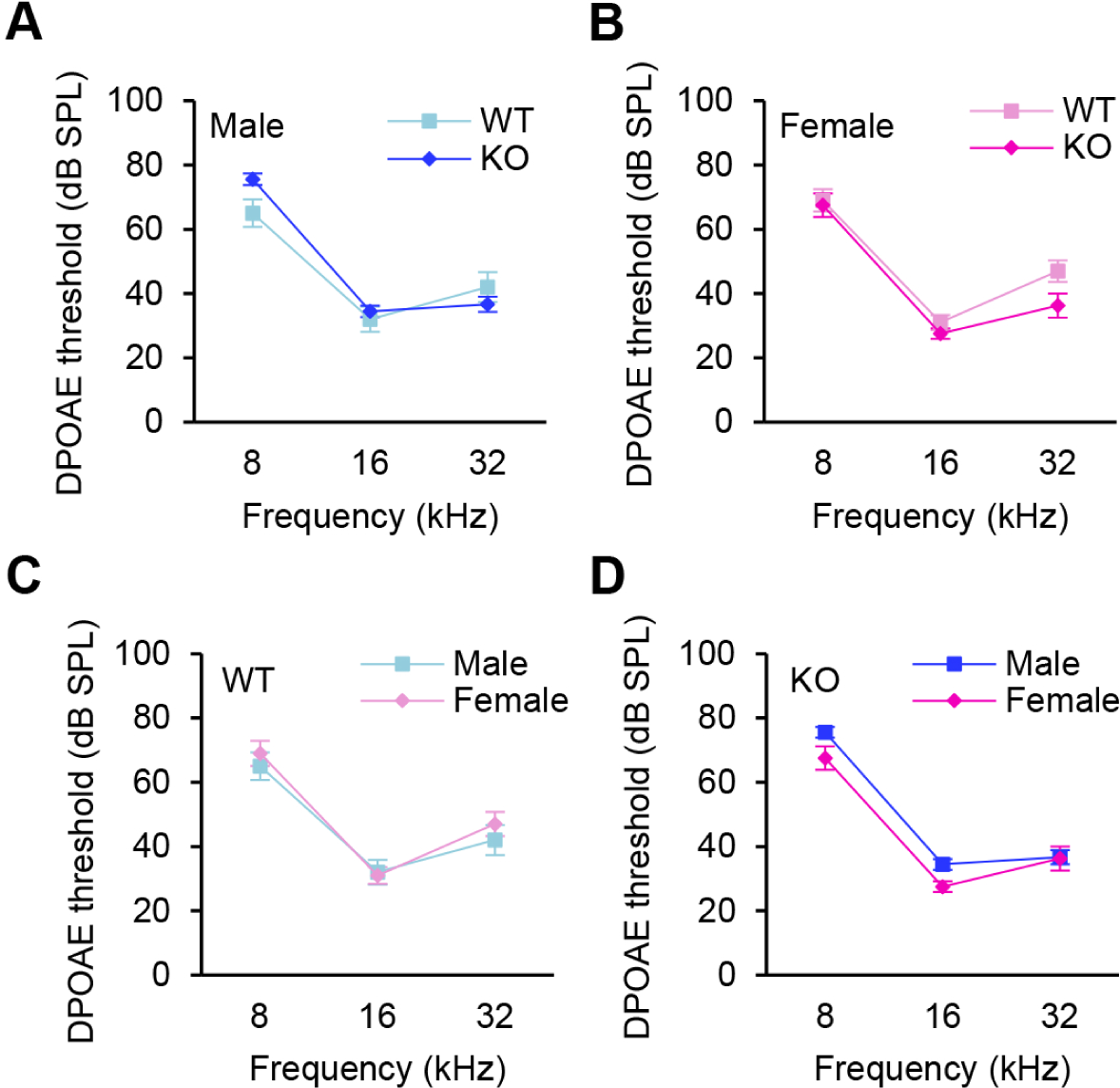
DPOAE threshold. (A-D) DPOAE thresholds were measured at 8, 16, and 32 kHz in WT and *Esr1* KO mice at 3 months of age; male WT (n=10), male KO (n=9), female WT (n=10), female KO (n=8). Data are shown as means ± SEM. Two-way ANOVA with Bonferroni’s multiple comparisons tests was performed. *p*<0.05.

### 3.3. Effects of Esr1 deficiency on rotarod balance performance

In rodents, physical activity and balance performance have traditionally been assessed by the rotarod test (Kim et al., 2023). To examine the effects of *Esr1* deficiency on balance performance, we measured rotarod balance performance (latency to fall) in WT and *Esr1* KO mice. There were no differences in rotarod balance performance between WT and *Esr1* KO males or WT and *Esr1* KO females, and there were no sex differences in rotarod balance performance in WT or *Esr1* KO mice (Fig. 8).

**Fig. 8.**
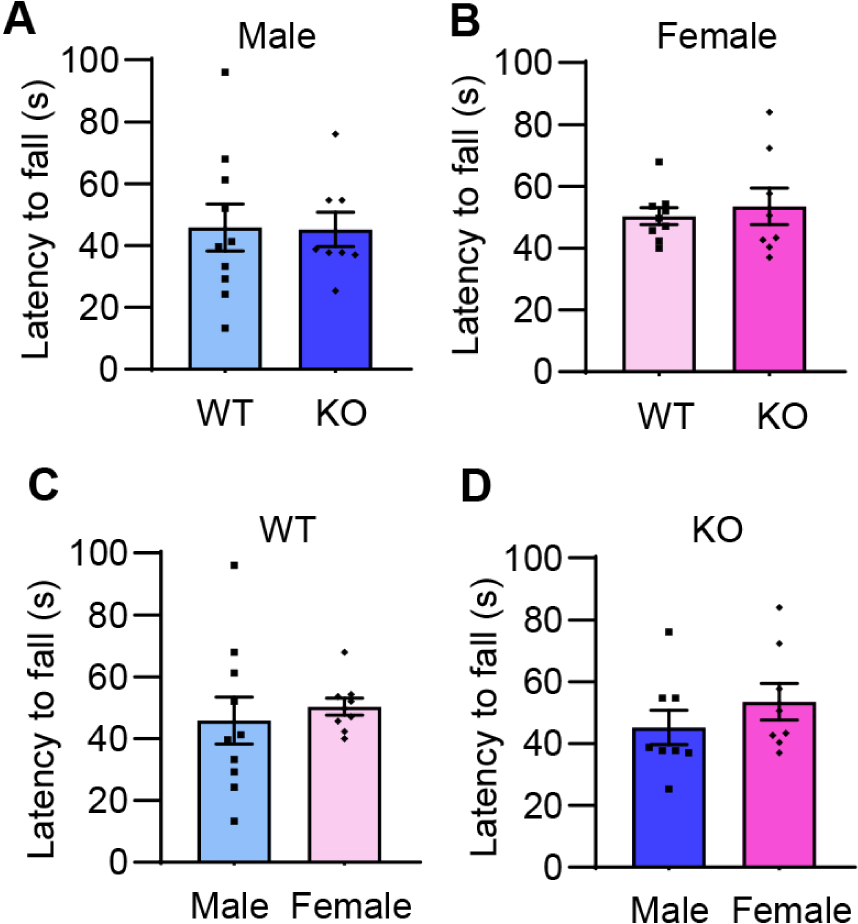
Rotarod balance performance. Rotarod performance was assessed by measuring latency to fall (s) in WT and *Esr1* KO mice at 3 months of age; male WT (n=10), male KO (n=9), female WT (n=10), female KO (n=8). Data are shown as means ± SEM. One-way ANOVA with Tukey’s multiple comparisons tests was performed. *p*<0.05.

## 4. Discussion

### 4.1. Effects of Esr1 deficiency on auditory function under normal physiological conditions

Our findings demonstrate that loss of *Esr1* does not affect ABR thresholds, ABR waves I-V amplitudes, ABR waves I-V latencies, DPOAE thresholds, or DPOAE amplitudes in normal-hearing young adult mice (CBA/CaJ background). These results are consistent with the previous report demonstrating that ABR thresholds from young (3 months old) *Esr1* KO mice on the 129/J-C57BL/6Ji background do not differ from WT mice (Meltser et al., 2008). In this study, the roles of ESR1, ESR2, and aromatase (CYP19A1 or ARO) were examined in response to auditory trauma using young mice deficient for *Esr1* (*Esr1* KO mice), *Esr2* (*Esr2* KO mice), or aromatase-deficient mice (*Ar*KO mice). The authors show that the mean pre-trauma ABR thresholds at 8, 12.5, 16, and 20 kHz from the *Esr1* KO, *Esr2* KO, and *Ar*KO mice do not differ from WT mice, indicating that loss of *Esr1*, *Esr2,* or *Aro* has no direct effect on basal hearing function in young mice. The authors also show that *Esr2* KO and *Ar*KO mice display increased susceptibility to acoustic trauma compared with WT, while *Esr1* KO mice did not show any differences in ABR threshold shifts after acoustic trauma compared to WT mice. Simonoska et al. (2009) also examined the role of ESR2 in auditory function using *Esr2* KO mice. At 3 months of age, there were no statistical differences in ABR thresholds at 6.3, 12.5, 20, and 40 kHz between WT and *Esr2* KO mice, and the inner ear morphology of *Esr2* KO mice appeared to be indistinguishable from WT mice. However, at 12 months of age, *Esr2* KO mice displayed profound hearing loss or near deafness, and the difference in ABR thresholds between the WT and *Esr2* KO mice was statistically significant for all tested frequencies. In the basal turns of the cochlea from the 12-month-old *Esr1* KO mice, no IHCs, OHCs, or spiral ganglion neurons were observed. In the basal turns of the cochlea from age-matched WT mice, there was also a loss of hair cells and spiral ganglion cells, but the loss was not as extensive as that observed in the *Esr2* KO mice. Taken together, our results and the previous reports demonstrate that loss of *Esr1* does not affect auditory function in young animals under normal physiological conditions. The previous studies also suggest that ESR2, but not ESR1, protects the auditory system from acoustic trauma or aging.

### 4.2. Effects of Esr1 deficiency on balance under normal physiological conditions

It is thought that estrogen affects the structure and function of the auditory system. However, the role of estrogens in vestibular function remains largely unknown. In rodents, balance performance and motor function have traditionally been assessed by the rotarod test (Kim et al., 2023). In the current study, our findings demonstrate that loss of *Esr1* does not affect rotarod balance performance in young mice. To our knowledge, this is the first report examining the role of ESR1 in balance performance in rodents. Simonoska et al. (2009) examined the expression of ESR1 and ESR2 in the inner ear of CBA/J mice, showing that ESR1 and ESR2 were co-expressed in the vestibular ganglion cells, vestibular dark cells, and endolymphatic sac. In 24-month-old mice, signal intensities of both ESR1 and ESR2 were significantly decreased in both sexes compared to young mice. These observations suggest that ESR1 has a role in normal vestibular organ function, but the loss of ESR1 is likely compensated by other estrogen receptors such as ESR2. In summary, our results show that loss of *Esr1* does not affect auditory or balance function in young animals under normal physiological conditions. Given that ESR1 is expressed in the inner hair cells, outer hair cells, spiral ganglion neurons, and stria vascularis (Charitidi et al., 2009; Motohashi et al., 2010; Stenberg et al., 1999), and in the vestibular ganglion cells, vestibular dark cells, and endolymphatic sac (Simonoska et al., 2009), further studies are needed to define the functions of ESR1 in the cochlea and vestibular organs not just in young, but also in middle-age and old mice using *Esr1* KO mice, *Esr2* KO mice, and mice deficient for both *Esr1* and *Esr2* on the same genetic background.

## Acknowledgments

This research was supported by the University of Florida’s University Scholars Program (Simms), R01 DC012552 (Someya) and R01 DC014437 (Someya) from the National Institute of Health and National Institute on Deafness and Communication Disorders, the Claude D. Pepper Older Americans Independence Centers at the University of Florida (P30 AG028740) from the National Institute of Health and National Institute on Aging, Evelyn F. McKnight Brain Research Foundation, and American Cancer Society (131062-RSG-17-171-01-DMC).

## Data availability

Data will be made available on request.

## Author contributions

**Shion Simms**: Writing – original draft, Writing – review & editing, performed genotyping and physiological experiments, analyzed and collected data, **Marcus Milani**: performed genotyping and physiological experiments, **Mi-Jung Kim**: performed physiological experiments, **Laura Infante**: analyzed physiological experiment data, **Ryan Husain**: performed physiological experiments, **Paul Cooke**: provided the *Esr1* mutant mouse line - Writing – review & editing, **Shinichi Someya**: Writing – original draft, Writing – review & editing, Supervision, Project administration, Funding acquisition.

